# Demonstrating the utility of flexible sequence queries against indexed short reads with FlexTyper

**DOI:** 10.1101/2020.03.02.973750

**Authors:** Phillip A. Richmond, Alice M. Kaye, Godfrain Jacques Kounkou, Tamar V. Av-Shalom, Wyeth W. Wasserman

**Author notes:** These authors contributed equally (Phillip A. Richmond, Alice M. Kaye). **Corresponding Author** Wyeth W. Wasserman < >.

## Abstract

Across the life sciences, processing next generation sequencing data commonly relies upon a computationally expensive process where reads are mapped onto a reference sequence. Prior to such processing, however, there is a vast amount of information that can be ascertained from the reads, potentially obviating the need for processing, or allowing optimized mapping approaches to be deployed. Here, we present a method termed FlexTyper which facilitates a “reverse mapping” approach in which high throughput sequence queries, in the form of k-mer searches, are run against indexed short-read datasets in order to extract useful information. This reverse mapping approach enables the rapid counting of target sequences of interest. We demonstrate FlexTyper’s utility for recovering depth of coverage, and accurate genotyping of SNP sites across the human genome. We show that genotyping unmapped reads can correctly inform a sample’s population, sex, and relatedness in a family setting. Detection of pathogen sequences within RNA-seq data was sensitive and accurate, performing comparably to existing methods, but with increased flexibility. We present two examples of ways in which this flexibility allows the analysis of genome features not well-represented in a linear reference. First, we analyze contigs from African genome sequencing studies, showing how they distribute across families from three distinct populations. Second, we show how gene-marking k-mers for the killer immune receptor locus allow allele detection in a region that is challenging for standard read mapping pipelines. The future adoption of the reverse mapping approach represented by FlexTyper will be enabled by more efficient methods for FM-index generation and biology-informed collections of reference queries. In the long-term, selection of population-specific references or weighting of edges in pan-population reference genome graphs will be possible using the FlexTyper approach. FlexTyper is available at https://github.com/wassermanlab/OpenFlexTyper.

**Author Summary:** In the past 15 years, next generation sequencing technology has revolutionized our capacity to process and analyze DNA sequencing data. From agriculture to medicine, this technology is enabling a deeper understanding of the blueprint of life. Next generation sequencing data is composed of short sequences of DNA, referred to as “reads”, which are often shorter than 200 base pairs making them many orders of magnitude smaller than the entirety of a human genome. Gaining insights from this data has typically leveraged a reference-guided mapping approach, where the reads are aligned to a reference genome and then post-processed to gain actionable information such as presence or absence of genomic sequence, or variation between the reference genome and the sequenced sample. Many experts in the field of genomics have concluded that selecting a single, linear reference genome for mapping reads against is limiting, and several current research endeavors are focused on exploring options for improved analysis methods to unlock the full utility of sequencing data. Among these improvements are the usage of sex-matched genomes, population-specific reference genomes, and emergent graph-based reference pan-genomes. However, advanced methods that use raw DNA sequencing data to inform the choice of reference genome and guide the alignment of reads to enriched reference genomes are needed. Here we develop a method termed FlexTyper, which creates a searchable index of the short read data and enables flexible, user-guided queries to provide valuable insights without the need for reference-guided mapping. We demonstrate the utility of our method by identifying sample ancestry and sex in human whole genome sequencing data, detecting viral pathogen reads in RNA-seq data, African-enriched genome regions absent from the global reference, and HLA alleles that are complex to discern using standard read mapping. We anticipate early adoption of FlexTyper within analysis pipelines as a pre-mapping component, and further envision the bioinformatics and genomics community will leverage the tool for creative uses of sequence queries from unmapped data.

## Introduction

Short-read DNA sequencing enables diverse molecular investigations across life science applications spanning from medicine to agriculture. Obtaining useful information from a data set of raw reads (short pieces of DNA read outs from the DNA sequencer) typically involves performing either *de novo* assembly, or mapping the read sequences against one or more reference genomes. Whether the focus is on quantification (e.g. observed gene expression in RNA sequencing data), or identifying sequence differences between a sample and a reference genome (e.g. genotyping), the availability of a curated reference genome has led to a large proportion of data analysis pipelines leveraging an indexed reference genome to perform efficient read mapping as a primary analysis component.

Recently a plethora of large-scale, population-specific sequencing projects have highlighted the numerous deficiencies and biases inherent to a single haploid reference (Ballouz, Dobin et al. 2019, Yang, Lee et al. 2019). Examples include the large amount of structural variation that exists between populations (Feuk, Carson et al. 2006, MacDonald, Ziman et al. 2014, Levy-Sakin, Pastor et al. 2019), the identification of unique sequences missing from the current reference genome (Sherman, Forman et al. 2019), and population specific difference in common genetic variants. Static linear reference genomes which do not capture these large differences between populations impose challenges for accurate genotyping, with implications in medicine and association studies (Ballouz, Dobin et al. 2019, Yang, Lee et al. 2019). Global efforts to enrich the linear reference genome have led to the development of graph based representations of pan-genomes, for a comprehensive review of current approaches see (Eizenga, Novak et al. 2020, Sherman and Salzberg 2020). As highlighted in an earlier review by (Paten, Novak et al. 2017), a key challenge in the future will be to determine the most appropriate reference genome(s), or path(s) through a graph pan-genome, to maximize genotyping performance. Knowledge regarding the genotypes of single nucleotide polymorphisms (SNPs) or other makers present in a read data set can be used to guide the choice of reference.

Currently, the approach of identifying SNP genotypes across the genome primarily involves computationally expensive reference-based read mapping and variant calling strategies (Nielsen, Paul et al. 2011). Recently published tools have highlighted the expanse of information that can be obtained from short read datasets. Inferring ancestry from specific, population-discriminating SNPs can be performed rapidly with Peddy, which uses fewer than 25,000 SNPs to identify ancestry through principal component analysis (Pedersen and Quinlan 2017). Somalier (Pedersen, Bhetariya et al.) avoids the final stage of variant calling and evaluates relatedness in aligned sequencing datasets. However, the accuracy of both of these tools is affected by the underlying alignment. Previous work has shown that it is possible to genotype predefined SNPs from unmapped sequence data, circumventing the read mapping and variant calling process (Shajii, Yorukoglu et al. 2016, Dolle, Liu et al. 2017, Sun and Medvedev 2019). Some approaches focus on k-mer (short sequences of length k) hashing and matching to predefined target k-mers to perform genotyping of known SNPs, as demonstrated in the VarGeno and LAVA frameworks (Shajii, Yorukoglu et al. 2016, Sun and Medvedev 2019). These approaches are fast, but rely upon indexes of k-mers extracted from the reference genome and SNP databases, thus reducing their flexibility for k-mers of different length and source. As we move into the era of precision medicine, avoiding inherent reference bias is crucial in obtaining accurate results. A separate approach is taken by Dolle et al., wherein the entire 1000 Genomes dataset is compressed into an FM-index and queried with k-mers spanning polymorphic sites, thus demonstrating the utility of scanning unmapped reads for predefined k-mers of interest. The “reverse mapping” highlighted in their approach was applied to aggregated data, but the concept can be extended to the analysis of individual genomes if implemented in a flexible way for diverse types of queries.

The reverse mapping approach switches the focus onto querying for sequences of interest within a read set, rather than a reference genome or database. This approach allows for a flexible exploration of the information content of the reads by allowing the read set to be queried for different parameters and across diverse sets of informative sequences. One example of this is within RNA sequencing (RNA-seq), where analysis of cancer RNA-seq datasets can reveal the presence of viral pathogens within patient data (Klijn, Durinck et al. 2015). Several tools have been developed to specifically detect these viral pathogens from sequencing data including viGEN (Bhuvaneshwar, Song et al. 2018) and VirTect (Xia, Liu et al. 2019). However, as with the tools mentioned earlier, they are hampered by a mapping procedure which first maps against the human reference genome and then subsequently maps against viral genome collections. Other methods, such as Centrifuge (Kim, Song et al. 2016) and Kraken2 (Wood, Lu et al. 2019), rely upon probabilistic or exact k-mer searches against large viral and bacterial databases. Both of these methods are powerful, but lack flexibility and rely upon phylogenetic relationships between target sequences. Specifically, they require the index for a search database to be recreated for different k-mer lengths or when additional target sequences are added to the database. Nevertheless, these tools are broadly used and thus serve as good comparators for efficacy, as they have both been demonstrated to have utility in detecting viral pathogens within cancer RNA-seq datasets by examining k-mer content. (https://www.sevenbridges.com/centrifuge/).

Combining the current drive to decrease our reliance upon linear reference genomes, and the wealth of demonstrated utility of reverse mapping approaches, we developed FlexTyper. FlexTyper is a computational framework which enables the flexible indexing and searching of raw next generation sequencing reads. We show example usage scenarios for FlexTyper by demonstrating the high accuracy of reference-free genotyping of SNPs in single samples, and the ability to identify foreign pathogen sequences within short-read datasets. We further explore the utility of FlexTyper within challenging genomic regions hampered by hyper-variability, and test its capacity to detect population-specific sequences missing from the reference genome. We hope the flexibility afforded by the framework underpinning FlexTyper will fuel the emerging trend away from the necessity for a static reference genome that currently lies at the heart of the majority of genomic analysis tools.

## Design and Implementation

### Overview of FlexTyper

Usage can be broken down into three steps: 1) query generation, 2) indexing the raw reads, and 3) querying against the FM-index (Figure 1). For query generation, we allow for both custom user query generation, as well as pre-constructed queries from useful databases, such as CytoScanHD array probe queries. Custom queries designed to capture genomic loci can be generated by pairing a user-provided VCF (format v4.3) with a reference genome fasta file. For the capture of potential pathogen sequences, we also allow query generation from one or more fasta files. The files produced from query generation are used as input for subsequent index query operations. The second step is the production of an FM-index from a set of short-read sequences in fastq, gzipped fastq or plain text format. There is the option to include the reverse complement of the reads within the index, however this increases the compute burden of indexing, without the same reduction in search times. The read set is concatenated using a sentinel character and passed as a single string into the indexing algorithm. The third step is the core FlexTyper search algorithm which takes the query input file, generates search k-mers, and scans the FM-index for matches. This step creates an output with matching format to the input query file, with appended counts of matching reads for each query. A detailed breakdown of these three components is described below.

**Fig 1.** Overview of FlexTyper. FlexTyper has three primary components: query generation, read indexing, and searching against the FM-index. Query generation includes the capacity to translate VCF files into query files given a reference genome file (e.g. Genome Fasta), or to directly create queries from fasta sequences including pathogen genome sequences. Modules VCF2Query.py and Fasta2Query.py facilitate this process. The second component involves creating an FM-index of the raw reads, after optional preprocessing steps. The third component searches the queries against the FM-index to produce output files with counts of query sequences within the query files.

### Query generation

FlexTyper supports flexible query generation giving users the capacity to query for any target sequence or allele within their read dataset. Query files can be generated from an input fasta and VCF file (VCF2Query.py), or directly from a fasta file (Fasta2Query.py). Potentially useful queries, including those presented here, are provided online and include all sites from the CytoScanHD chromosomal microarray, and ancestry discriminating sites (Pedersen and Quinlan 2017). These predefined query sets are available through git-lfs in the online FlexTyper Github repository (https://github.com/wassermanlab/OpenFlexTyper). If users wish to directly query a short-read dataset with a set of predetermined k-mers, we provide a separate function, ksearch, that will directly search for k-mers from a given text file within an indexed read set.

### FM-index creation

Generating the FM-index from short-read sequencing datafiles is performed in two steps: preprocessing and indexing. The focus of our work is not on the algorithms used to construct the FM-index, and hence we use two existing utilities to generate a compatible FM-index for FlexTyper. The toolkit Seqtk is used for reformatting the input read files by removing quality scores and non-sequence information to create a sequence-only fasta format. The output fasta file is processed using the SDSL-Lite library to generate the FM-index. SDSL builds a suffix array that is used to generate the BWT of the input string, which is then compressed using a wavelet tree and subsampled. The resulting compressed suffix array is streamed to a binary index file. As the memory requirements for indexing large files can be burdensome, we support an option to split the input file and index each chunk of reads independently. Downstream search operations support the use of multiple indexes.

### Query against FM-index

Querying the FM-index for user selected sequences can be conceptually divided into four steps: 1) k-mer generation; 2) k-mer filtering; 3) k-mer searching; and 4) result collation (Figure 2). There are two primary methods of k-mer generation for a query; a centered search where the middle position of the query is included in all k-mers, and a sliding search which starts at one end of the query and uses a sliding window approach to generate the k-mers (Figure 2). Centered search can be used for genotyping or estimating coverage over a single position, and the sliding search can be used to count reads which match to any part of a query sequence. All parameters for the search are specified in the settings.ini file, with a small number of key parameters able to be overridden directly from the command line. After filtration, the k-mers are searched for within the FM-index using C++ multithreading and asynchronous programming, using either a single thread on a single index, multiple threads on a single index, a single thread on multiple indexes, or multiple threads on multiple indexes (Figure 2). Importantly, asynchronous programming allows the number of threads used during searching to be increased beyond the number of available CPUs. The output from this search process is a collated results map containing the positions of each k-mer within the FM-index. These positions are translated to read IDs, and finally collapsed into query counts using the k-mer-query map. Importantly, if multiple k-mers from the same query hit the same read, they are recorded as a single count at the query level. For cases of multiple indexes being searched in parallel, the k-mer searching is performed independently for each index, and then the search results from all indexes are merged and reconciled to produce a final query count table. For a detailed explanation of the effects of key parameters, please see Supplementary Information, and our documentation on Github pages (https://wassermanlab.github.io/OpenFlexTyper/)

**Figure 2.**
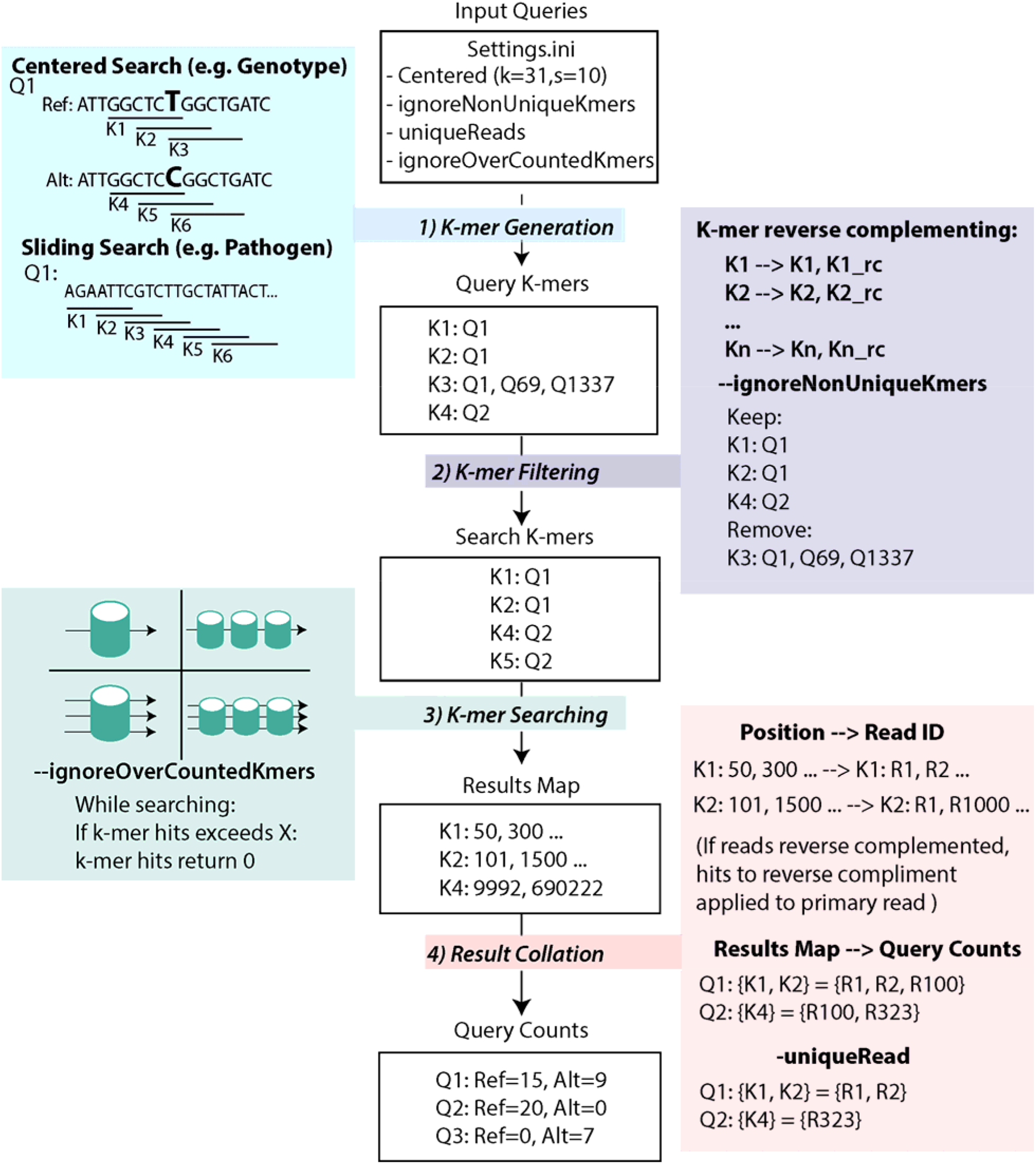
Query Search Workflow. Workflow for query search against the FM-index, starting with input queries and settings defined in Settings.ini file. In this figure, the example shows a centered search with ignoreNonUniqueKmers enabled. 1) K-mer generation has two modes centered search and sliding search. For a centered search, the position of interest lies in the middle of the query, and k-mers are designed to overlap that central position with defined length (k) and step (s). 2) If the ignore-duplicates option is set, k-mers collated from the query set are filtered to remove any k-mers which were found in multiple query sequences. 3) The filtered k-mers are then searched for within a single FM-index (left two panels) or multiple indexes (right two panels) of the read set. This can be done using single (top two panels) or multiple (bottom two panels) threads. 4) The results corresponding to a position within the FM-index are then translated back into reads, with hits on reverse complement reads assigned to the primary read, and collapsed into a set for each query. The final counts are reported per query.

### Post-processing of results into downstream formats

The output tables from the search process for genotyping can be translated into useful formats for downstream analysis using the fmformatter scripts (https://github.com/wassermanlab/OpenFlexTyper/tree/master/fmformater). Currently, there is the capacity to output genotype calls in VCF, 23andMe, or Ancestry.com format. Genotype calls are derived here using a basic approach which assigns genotypes given a minimum read count parameter as follows:

Alt < minCount && Ref > minCount: Homozygous reference, 0/0
Alt > minCount && Ref > minCount: Heterozygous alternate, 0/1
Alt > minCount && Ref < minCount: Homozygous alternate, 1/1

For searches which do not pertain to genotyping, the output tab-separated files can be used as count tables for observed query sequences.

## Results

### Observations about FlexTyper system requirements and performance

The generation of a full text index of the reads is a key step and we are able to generate indexes of human whole genome sequencing reads with ~800 million reads utilizing less than 150Gb of RAM on a single compute node within a higher performance compute (HPC) cluster. (Supplemental Table S1). Although read indexing is slower than a traditional alignment, sorting, and variant calling pipeline, FlexTyper can index whole genome sequencing samples in roughly 24 hours (Supplemental Table S1). The flexibility for creating sub-indexes allows the user to adjust parameters to fit their system, accommodating most modern HPC architectures. In comparison to the tools that utilize prebuilt indexes, such as BWA-MEM, or generate probabilistic indexes, such as Kraken2, FlexTyper is significantly slower, however, the non-approximate full text implementation allows the read set to be queried for diverse sequences, across the full parameter space, without reindexing unlike Kraken2. The index of a high depth paired-end (2×250bp) WGS read set for sample HG002 uses only 155 Gb RAM, and with no information loss in the read sequences, it is not necessary to retain the original fastq files once indexed. While FlexTyper does remove quality scores, the tuning of k-mer length and step size allows counting of a read even in the presence of errors, and filtering the fastq file to remove low-quality/high-error reads can be done prior to read indexing. The complex interplay of the different search parameters makes generalized performance statements challenging, so to inform the user of FlexTyper’s use-case performance we provide runtime metrics and read recovery for a variety of different search settings and scenarios in the following sections.

### Testing for the presence of pathogen sequences in RNA-seq

To demonstrate the capacity of FlexTyper to detect pathogens from RNA-sequencing data, we generated synthetic reads from four relevant viral genomes including Epstein-Barr virus (EBV), Human Immunodeficiency virus type 1 (HIV-1), and two Human Papilloma virus strains 68b (HPV FR751039) and 70 (HPV U21941) (Supplemental Methods). We first examined the impact of various FlexTyper parameters on the recovery rate of pure, simulated read sets for each of the four viruses and one human blood RNA-seq dataset from the Genome England project (https://www.ebi.ac.uk/arrayexpress/experiments/E-MTAB-6523/samples/) (Table S2). Importantly, varying the parameters *k* (length of the k-mer search substring) and *s* (step-size) change the specificity and sensitivity of read recovery. When *k* is set to 15 (a short k-mer), there were roughly 1 million off-target hits to the viral genomes for the pure human RNA-seq file, and the increased search space leads to increased run times (Table S2). Next, we explored the effect of two uniqueness settings, the first for k-mers, (ignoreNonUniqueKmers), where a given k-mer cannot map across multiple queries, and secondly for reads, (uniqueReads), where a read cannot be counted across multiple queries. We provide both parameters to the users, as there may be instances where reads are allowed to be counted across queries, but the k-mers must be independent. By exploring these parameters, we show that all simulated reads can be recovered with parameters of k=30 and s=5, and off-target assignment can be controlled with the uniqueness settings. For more details about the parameter explanations or the impact of parameterizations on read recovery, see Supplemental Information and Table S2.

Next, we simulated mock patients infected by the four viruses to examine the detection capacity of FlexTyper with respect to two established methods, Centrifuge and Kraken2. Simulated reads from each of the four viruses were spiked-in at various concentrations (read counts) within five different human RNA-seq datasets from the Genome England project (https://www.personalgenomes.org.uk/data/) (Supplemental Methods). Using the optimized parameters derived above (k=30, s=5, uniqueRead, and uniqueK), we are able to detect each virus in the patient samples even at low concentrations (Figure 3). We also searched these mock patient samples using an expanded set of parameters, and see the expected changes in sensitivity as the uniqueness and k parameters are altered (Table S3). For more direct comparison to Centrifuge and Kraken2 we ran FlexTyper in the paired-read mode, and show that we are able to detect the pathogen sequences with high sensitivity and specificity (Figure 3). This is true even for a sample (Patient 5) which had a 1x concentration–roughly 50-1150 reads depending on genome size–of each viral genome spiked in. For Patient 4, where the level of EBV spiked in is over 1 million reads, the undercounting stems from our limit on maximum occurrence, which we set to 2000, and adjusting to a maximum occurrence of 10,000 increases the count to 1,144,990. For the HPV viruses (U21941 and FR751039), we are able to detect the levels of both strains, and observe a slight over-counting of FR751039 in Patient 2, possibly from a high count of spiked in U21941 reads. We compared our approach with Centrifuge and Kraken2, which match reads based on k-mers mapped against a comprehensive indexed viral and bacterial database, and tabulate matches at the read pair level. Centrifuge works with unmapped short-read sequencing data by performing read-length (k=150) k-mer searches against a database of viral and bacterial genomes, and hence was the least sensitive method due to the limitations of full length k-mer queries (Kim, Song et al. 2016). Kraken2, which uses a minimizer for approximate matching, can search for shorter k-mers (default k=31), leading to increased sensitivity over the Centrifuge method. Both Centrifuge and Kraken2 achieve accurate results for the HIV-1 and EBV samples, but were only able to detect 5-10% of the reads for the two HPV samples, U21941 and FR751039, even when aggregating at the family level (Papillomaviridae) (Figure 3). Initially, we hypothesized that this was due to off-target mapping to another genome, perhaps the human genome, within their comprehensive database. However, after testing a set of 5200 pure U21941 or FR751039 reads with Kraken2, we were only able to recover 136 reads and 279 reads respectively, even when considering reads assigned to the viral kingdom level (Supplemental Methods). This limitation can be overcome with FlexTyper, which enables the user to define the relevant pathogens to search for, along with the ability to repeat searches across different k-mer lengths, without the need for re-indexing a complex bacterial or viral database. While Kraken2 and Centrifuge are powerful and comprehensive metagenomic classifiers, which allow for increased breadth and classification across a phylogenetic tree, there may be cases where specific pathogen queries of interest require high sensitivity. We believe FlexTyper can serve as an option in these scenarios.

**Figure 3.**
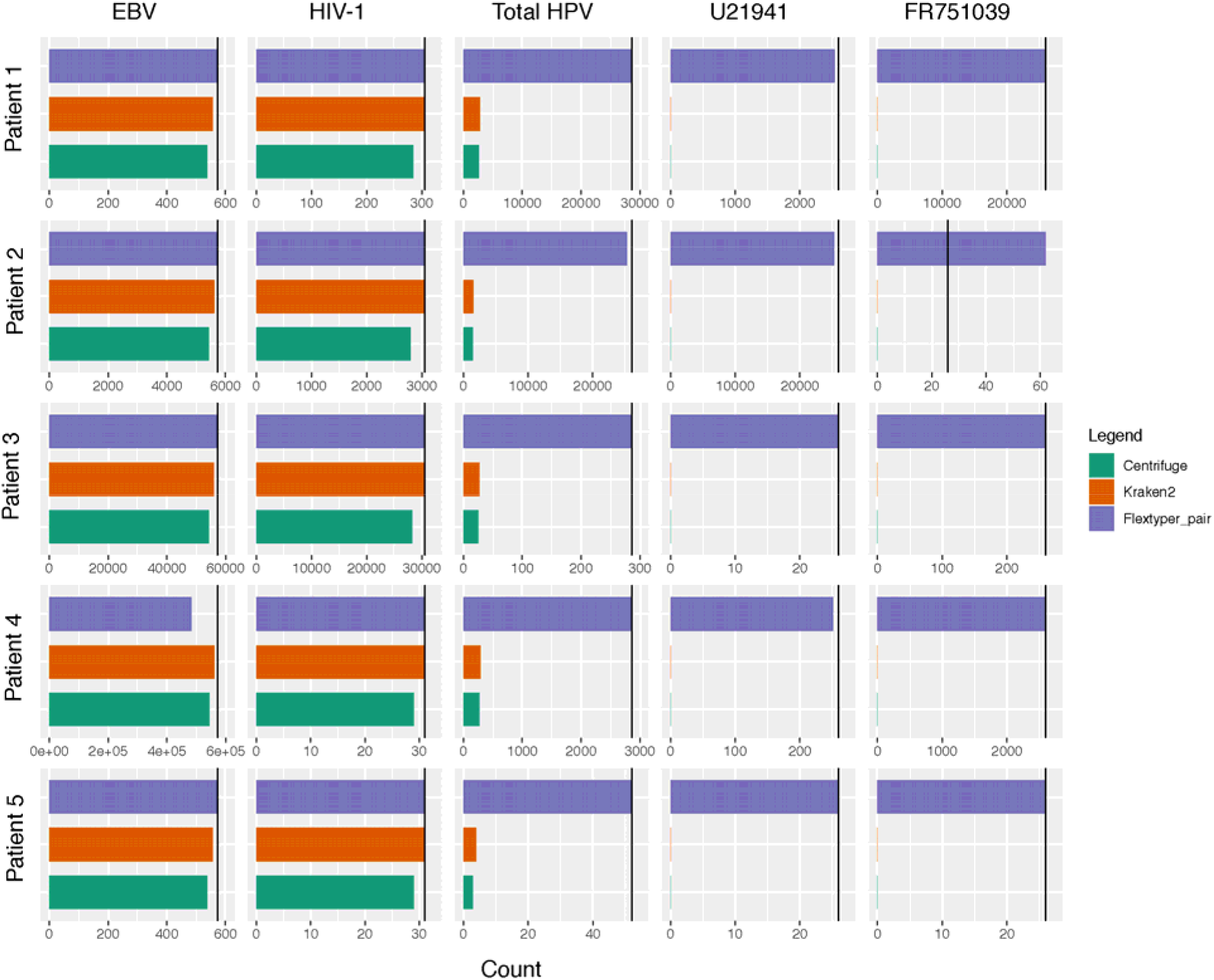
Mixed Viral Analysis. Detection of pathogen sequences in five synthetic patient RNA-seq datasets (Patient 1-5; rows), each with different levels of spiked-in viruses (EBV, HIV-1, U21941, and FR751039; columns), expected values shown as black vertical bars. As Centrifuge and Kraken2 are unable to delineate between the two HPV substrains (U21941 and FR751039), a combined count at the HPV level is tabulated.

### Genomic coverage and genotype detection within human WGS data

Knowing whether a given k-mer is present or absent from a human WGS datafile (in this instance Illumina short-read, paired-end data) can have utility for estimating the depth of coverage for a target region and genotyping SNPs. FlexTyper has the capacity to compute depth of coverage or genotype SNPs from WGS data for both predefined and user-supplied loci. We demonstrated this capacity for genomic sites using the probe sequences from the CytoScanHD microarray, as well as a subset of previously collated population discriminating SNPs (Pedersen and Quinlan 2017). Using these loci, we created query files with a reference and alternate query sequence centered on the biallelic site (Supplemental Methods).

We first sought to test the read recovery capacity of FlexTyper compared to an alignment based method which we call BamCoverage. The BamCoverage method involves mapping the reads to the reference genome, and then extracting per-base read coverage over a specific reference coordinate. BamCoverage utilizes the pysam package to extract read pileup over positions defined by the FlexTyper input query file (Supplemental Methods). Using the CytoScanHD SNP set, we found a high concordance between the read counts from FlexTyper and the depth of coverage from aligned reads (Figure 4A). FlexTyper was run with parameters of k=31, s=10, max occurrence 200, and the requirement of unique k-mers between queries. The vast majority, 98.33% (784,297/797,653), of sites differed by less than 10 between FlexTyper and BamCoverage, with a Spearman correlation of 0.86 (Figure 4B). The discrepancy in counts is similar for both reference and alternate alleles, which is important since most genotyping models assume relative contributions of observed alleles for genotype calling. There were 10,397 sites with a delta, (Δ = FlexTyper - BamCoverage), greater than 10, and 1,469 sites with a delta greater than 100 (Figure 4C). We manually investigated a few of these sites which were overcounted by FlexTyper by more than 100 and found that they are being overcounted due to k-mers mapping to multiple possible locations. Comparing these over-counted hits with delta greater than 100 to previously defined repeat regions shows that 1444/1469 or 98.3% of the overcounted sites overlapped with predefined repeats (Trost, Walker et al. 2018). The uniqueness of k-mers is important for accurate read counting, thus it is recommended to filter query sequences within such regions when using FlexTyper for genotyping or depth profiling. Lastly, by examining the recovery of reads across the chromosome between FlexTyper and the BamCoverage approach, we observe uniform recovery across the breadth of the chromosome (Figure 4D). This is important for copy number variant calling applications, as they rely upon contiguous readouts of genomic sequence coverage.

**Figure 4.**
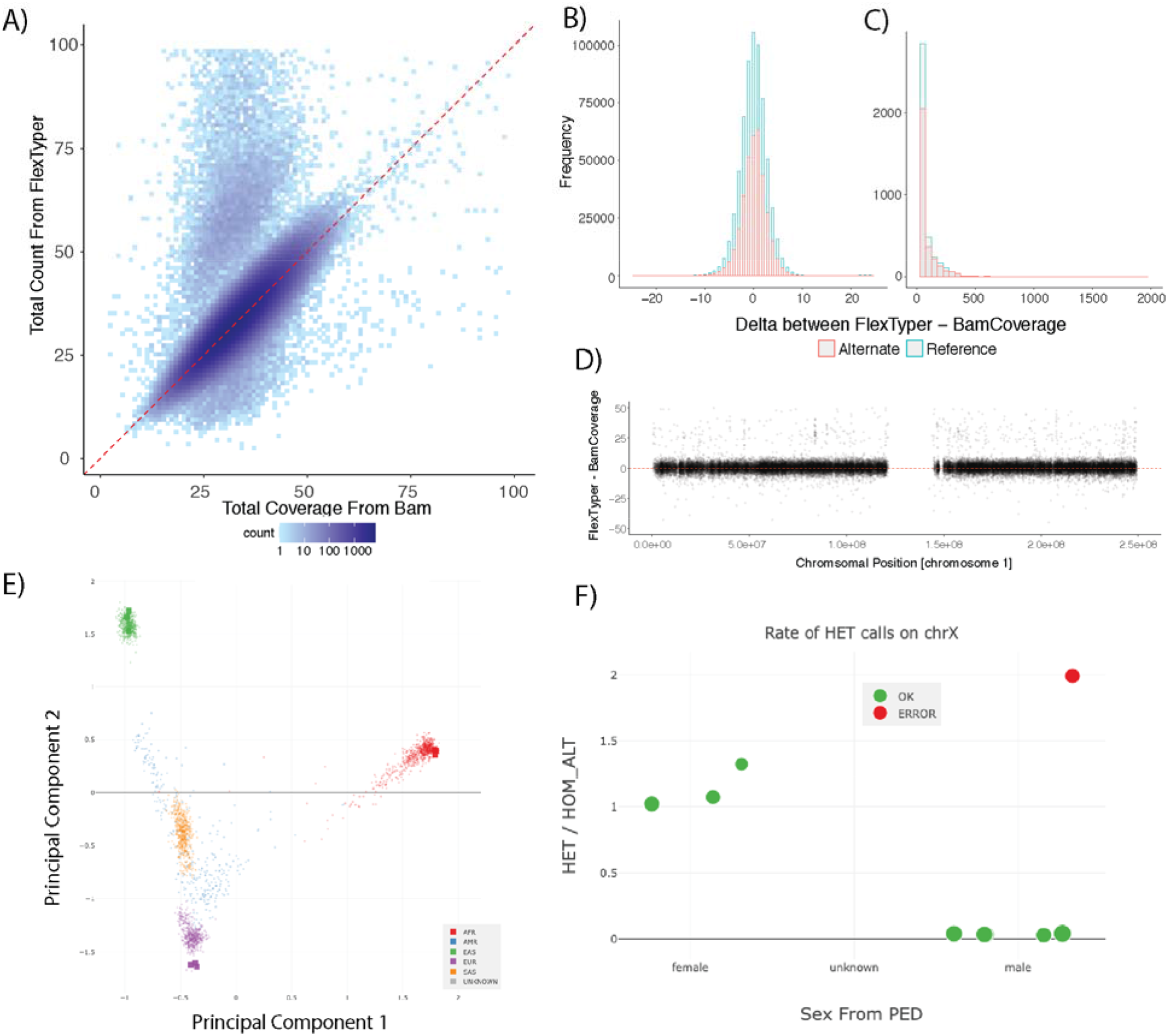
WGS Genotyping using FlexTyper. A) FlexTyper read count compared to the total coverage from BAM file over SNP sites represented on the CytoScanHD microarray. B) Histogram showing the delta, (Δ = FlexTyper - BamCoverage), in read count for both the alternate (red) and reference (blue) alleles. C) Histogram of the same delta as B) but with an extended axis from 100-2000, showing the frequency of over-counting for sites using FlexTyper. D) Scatter plot showing the delta (Δ = FlexTyper - BamCoverage) on the y-axis, plotted across chromosome 1 on the x-axis. E) Principal component analysis showing projection of FlexTyper-derived SNP genotypes from nine individuals of Asian (green), African (red) and European (purple) ancestry. Squares denote FlexTyper genotypes, points denote existing data from the 1000 Genomes project provided by Peddy. F) Sex-typing for these Polaris samples showing the ratio of heterozygous to homozygous sites on the X chromosome (y-axis) for individuals for the defined sexes as male (right) and female (left). Each individual is labeled as green (correctly sex-labeled) or red (incorrectly labeled).

Next, we investigated whether FlexTyper can accurately recover genotypes at the SNP sites profiled from the chromosomal microarray. The genotyping approach we use leverages a minimum count from the reference and alternate allele to assign heterozygous, homozygous alternate, or homozygous reference genotypes (Supplemental Methods). We applied this basic genotyping algorithm to both FlexTyper and BamCoverage counts to produce a VCF file. These genotypes were compared to an alternate pipeline which uses reference-based mapping and sophisticated variant calling using BWA-MEM and DeepVariant (Poplin, Chang et al. 2018) (Supplemental Methods). For the CytoScanHD microarray, all three methods report the same genotype for 99.4% (792,805/797,653) of the SNP sites. We further investigated the discordant genotypes to see if we can explain why there is a disagreement between FlexTyper and the other two methods. First, we compared these genotypes to the repeats defined above and see that 77.6% (2,980/3,838) of the discordant genotypes overlap with predefined repeat loci. Additionally, we implemented a flag for offending k-mers, to signal when a k-mer was non-unique or surpassed the max occurrence (maxOcc) parameter. There were a total of 685 genotypes with offending k-mers, of which 355 (8.6% of the 3,838) overlap with the discordant genotypes unique to FlexTyper. We further demonstrate the accuracy of FlexTyper-derived genotypes by indexing nine WGS samples from the Polaris project representing diverse populations including three African, three Southeast Asian, and three European individuals (Chen, Krusche et al.). After indexing, we queried the samples for population discriminating sites and then genotyped the output table to produce a VCF file. The output VCFs were then used within the Peddy tool, and a principal component analysis was performed to predict the ancestry of the samples (Pedersen and Quinlan 2017). In all nine cases the population was correctly determined, as well as the relatedness inference for the three trios (Figure 4E, Figure S1). Interestingly, we observed a discrepancy between the listed sex for the child of the European trio, individual HG01683, and the inferred sex from FlexTyper and Peddy (Figure 4F). We followed up on this observation and revealed that the individual is not an XY male, but rather an XXY individual, and communication was made that resulted in the relabeling of the individual within the online repository. The analysis time for extracting the queries from the indexed reads for all WGS analysis can be found in Table S4, highlighting that especially for informative subsets of queries, such as population discriminating sites, genotypes can be recovered accurately and quickly (~10-15 minutes). Taken together, FlexTyper has the capacity to provide accurate counts of observed reads matching two alleles over informative SNP sites, with relevant utilities such as copy number estimation, sample identification, ancestry typing, and sex identification.

### Exploring creative uses of FlexTyper

To demonstrate the flexible utility of our k-mer-based searching method, we explored regions of the genome which are challenging for read mapping and downstream analysis when represented within haploid, linear reference genomes. The two areas we chose to focus on include contigs derived from a population but not present in the reference genome, and hypervariable and homologous regions, where linear representations are known to perform poorly.

The contigs we chose to process include the “non-reference” contigs from a recent African pan-genome publication (Sherman, Forman et al. 2019). These contigs, which were assembled from non-mapped reads, collectively contain nearly 300 megabases of DNA, represented by ~125,000 contigs. We created queries of these contigs, and then searched them using the nine samples from our ancestry WGS experiment using parameters of k=50, s=5, and uniqueRead=true. Unexpectedly, when we queried the African contigs across the family trio samples from three populations (EAS, EUR, and AFR), we observed similar contig coverage across all individuals (Figure 5A). Next, we sought to identify discriminating contigs within the ~125,000 non-reference contigs by filtering for those which consistently appear in one population (>10 counts in mother, father, and child), but had low coverage in the other two populations (<5 counts). Applying this to the three groups, we identified a set of discriminating contigs (Figure 5B). The African trio had 151 unique contigs, the East Asian trio had 60 unique contigs, and only four contigs were unique to the European trio. In total, the analysis indicates that the African-derived contigs are widespread across populations. Two limitations of our analysis include: 1) FlexTyper was run in unique mode, so reads mapping across highly similar contigs are discounted, and 2) FlexTyper does not account for local genome context, so it is possible that some of the contigs are unique not due to specific sequence, but due to their placement in the genome (e.g. structural variants). This application highlights the potential of FlexTyper in filtering and querying for contigs unique to a subpopulation.

**Figure 5.**
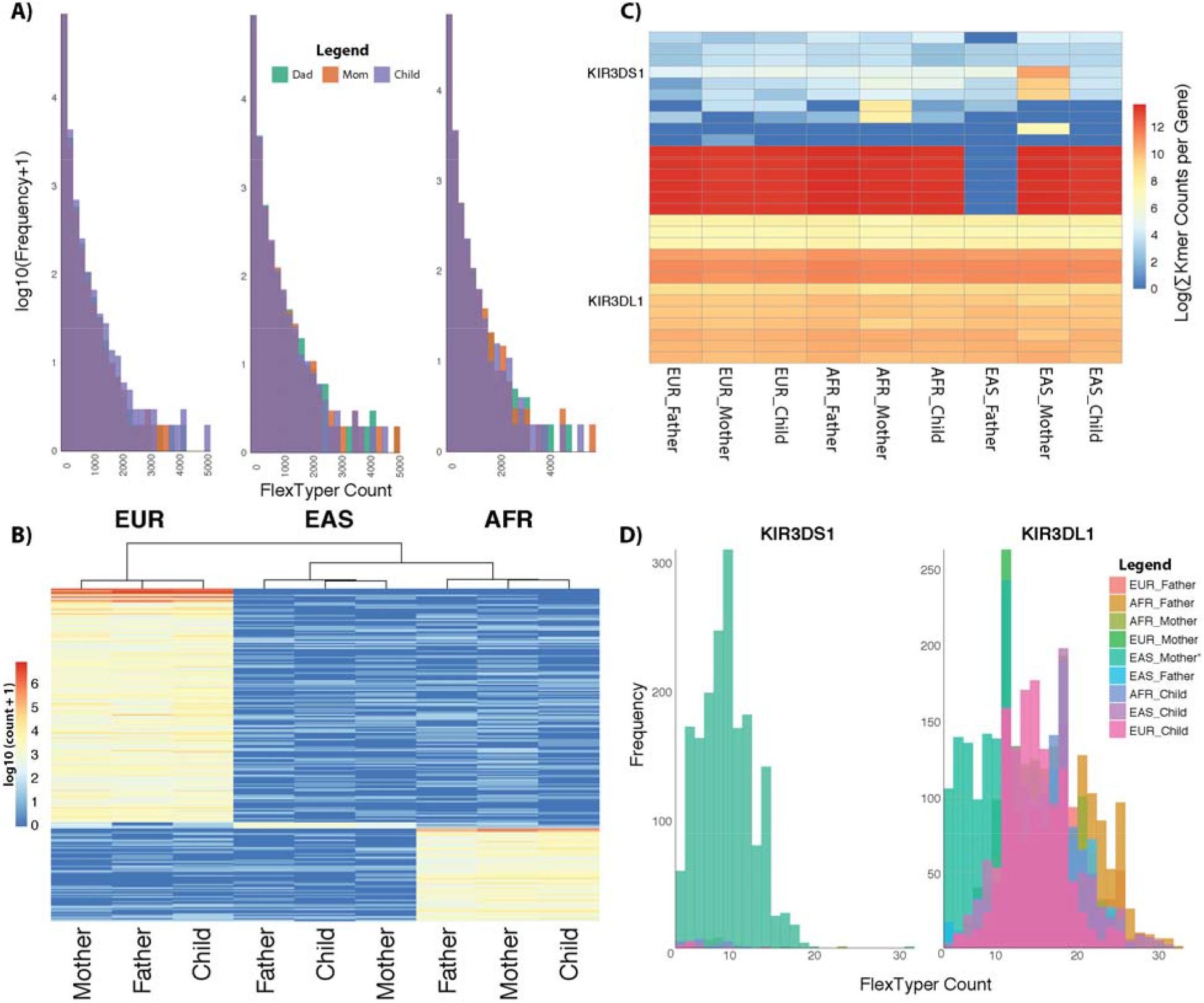
Explorative uses of FlexTyper. Two examples of the creative uses of FlexTyper within challenging regions. A) Histograms for counts of the African contigs overlaid for father (green), mother (orange) and child (purple) split by population (left to right EUR, EAS, AFR). B) Heatmap in log-scale for population-specific contigs, clustered by sample similarity (columns) and contig count similarity (rows). C) Heatmap showing the log10 transform of the sum of k-mer counts per gene, with genes as rows, and samples as columns. The two alleles, KIR3DS1 and KIR3DL1 are labeled as rows on the left side. D) Overlayed histograms for the 9 samples, showing the frequency of the FlexTyper k-mer count for the KIR3DS1 (left) and KIR3DL1 (right) alleles.

Recent tailored approaches to genotyping challenging genomic regions, which are difficult due to their hypervariability in the population and/or high sequence similarity between homologous genes, utilize unique k-mer counts to distinguish between alleles present in a sample (Roe and Kuang, Shen and Kidd 2020). As FlexTyper has the capacity to rapidly query k-mers and generate unique k-mers across input queries, we decided to test FlexTyper’s utility in distinguishing between samples for a locus known to be challenging: the killer-cell immune receptor (KIR) locus. We downloaded a curated set of gene-distinguishing k-mers for this locus which have been used, with the k-mer counting tool KMC3, to identify the presence-absence of the 28 genes/alleles in the KIR locus (Roe and Kuang). Using the FlexTyper function ksearch, we searched the nine WGS samples for this set of k-mers, and then tallied the k-mers with >3 counts per gene (Figure 5C). While we did not see many differences at the family level with this set, we did observe an outlier sample: the mother in the East Asian trio. For the KIR3DS1 gene, she had several high-counting k-mers which were absent in the other samples. KIR3DS1 is an alternate haplotype of the KIR3DL1 gene in the canonical reference genome, and is represented in the GRCh38 reference on an ALT contig (chr19_KI270887v1_alt). By plotting a histogram of the counts for the nine individuals over both the KIR3DS1 and KIR3DL1 genes, we observed that the mother of the East Asian trio is the only sample in this set with k-mer coverage over KIR3DS1, and consequently has reduced coverage of the KIR3DL1 gene (Figure 5D,E). Taken together, this suggests that the mother is heterozygous for the KIR3DL1 and KIR3DS1 alleles, while the rest of the samples in this set are homozygous for the KIR3DL1 allele. This observation is enabled by the careful selection of k-mers by Roe et al., and is a demonstration of how FlexTyper can utilize user curated k-mers for genotyping within a challenging locus.

## Discussion

Here we presented FlexTyper, a user-friendly tool which enables exploratory analysis of short read datasets without the need to perform reference guided alignment. Our framework allows the user to generate custom queries, or to directly search from a list of k-mers. This gives the user complete flexibility to tailor the search inputs and parameters to the problem at hand. We demonstrated three common applications: depth of coverage analysis, accurate SNP genotyping, and sensitive detection of pathogen sequences. We then showcased the potential for FlexTyper to extract useful information from complex, hypervariable, or non-reference genomic sequences. FlexTyper was designed with user simplicity in mind, but without comprising the breadth of potential applications, and hence the tool is available for the creative use of genomics researchers.

The rapid and accurate recovery of read depth enables innovative usage of FlexTyper in the space of copy number variant profiling. We demonstrated that we could reproduce the depth of coverage of a genomic region without the need for reference-based mapping. As microarrays begin to be replaced by genome sequencing assays, we envision that FlexTyper could be extended to reproduce microarray-style outputs that are established in clinical labs. Further, we show that when genomic queries with counts higher than the expectation arise, these events correspond to repetitive genomic sequences. As such, FlexTyper may not only enable the recovery of read depth in an accurate manner, but it can also inform the quality of a sequence query as a “unique probe” for assessing genomic copy number.

The genotyping case study highlights how pre-alignment analysis of genome sequence data can provide rapid insights into the properties of a sample. SNP genotyping was accurate across the genome, allowing rapid identification of sample ancestry, sample relatedness in the trio setting, and sample sex typing using Peddy (Pedersen and Quinlan 2017). Interestingly, applying Peddy to the output of FlexTyper for open source trio data from the Polaris project revealed a mislabeling of the sex for individual HG01683, which was reported and subsequently amended in the online data repository (https://github.com/Illumina/Polaris/wiki/HiSeqX-Kids-Cohort). Since ancestry and sex information can inform choices in downstream data processing, identifying these discrepancies between labeled sex and inferred sex in a data-driven manner is a critical step of pre-alignment informatics. For instance, mapping against the sample-matched sex chromosomes has been shown to improve performance (Olney, Brotman et al., Webster, Couse et al. 2019). As such, using FlexTyper, in combination with Peddy, on diverse datasets prior to reference-guided read alignment will lead to improved results from mapping-based pipelines.

There is increased recognition of the important of pathogen detection. In both cancer profiling (Klijn, Durinck et al. 2015) and public health studies (Gardy, Loman et al. 2015), rapid determination of the presence of pathogen sequences could obviate the need for full reference mapping. Some existing tools designed for viral detection in sequencing data rely upon pre-indexed databases of viral and bacterial sequences, sometimes including a phylogenetic relationship between genomes within the index (Kim, Song et al. 2016, Wood, Lu et al. 2019, Xia, Liu et al. 2019). Two approaches, Centrifuge and Kraken2, have been applied to cancer genomes to confirm the presence of viral pathogens, including Human papilloma virus (HPV). We demonstrated that our approach compares favorably to Centrifuge, with a more sensitive detection level, due to the ability to search for k-mers shorter than the read length and the advantage of fine-tuned control over the searchable database. Comparing FlexTyper to Kraken2, which doesn’t rely upon full read length queries, detection of the spiked-in pathogen sequences was as good or better than Kraken2, with improved performance for detecting the HPV-derived reads. Interestingly, both Kraken2 and Centrifuge had difficulties in detecting HPV reads, both within mixed-virus and pure viral read sets. Here we only searched for viral pathogens of interest, although other specific pathogen queries could be performed, such as the presence of antibiotic resistance genes within a patient RNA-seq sample.

As the research community begins to move away from a single haploid reference towards richer pan-genome representations, we anticipate that more diverse and creative uses for FlexTyper’s ‘reverse mapping’ approach will emerge. During our continued exploration of FlexTyper’s potential, we have identified a few possible applications. We focused on regions of the genome which are challenging for traditional linear reference approaches, including a set of sequences not present in the reference genome, and a highly polymorphic region with homologous genes. Using the set of contigs assembled from the African Pan-genome project, we applied FlexTyper and observed similar sequence coverage over these contigs across three families from European, East Asian, and African ancestry. We further filtered this set of contigs and identified a limited set of discriminating contigs, highlighting another relevant use case for FlexTyper. Beyond non-reference contig searching, we explore the utility of FlexTyper in genotyping the polymorphic and homologous genes within the KIR locus. We use a curated set of k-mers from a work by Roe et al, and identify an alternate haplotype (namely KIR3DS1, present in the ALT contigs of GRCh38) for the KIR3DL1 gene in one of the nine individuals. This genotyping demonstration with a curated set of k-mers highlights the potential for FlexTyper to be adopted by other specialized methods tailored for challenging genomic regions.

The full breadth of possible applications of FlexTyper and its reverse mapping approach has yet to be discovered, but we have highlighted multiple potential avenues here. For WGS read data sets, it is feasible to genotype complex structural variants by searching for sequences overlapping breakpoints, such as those observed in a subpopulation, or events recurrently found in cancer (Sudmant, Rausch et al. 2015, Li, Roberts et al. 2020). Within RNA-seq data, querying for exon-exon splice junctions in a rapid manner can allow isoform quantification, as has been previously demonstrated (Patro, Mount et al. 2014, Bray, Pimentel et al. 2016). Further, a recent report showed the utility of k-mer-counting methods in resolving copy number variants within paralogous loci and genes (Shen, Shen et al. 2020). Another group showed the advantage of examining depth of coverage at specific sites across the paralogous genes in Spinal Muscular Atrophy (Chen, Sanchis-Juan et al. 2020) As FlexTyper is well suited for specific sequence recovery operations, scanning with preselected query sequences such as defined by these studies can enable rapid detection (Chen, Sanchis-Juan et al. 2020). All of these proposed applications help tackle challenges which are currently a burden for traditional reference-based mapping approaches.

We focused this report on the expansive utility of querying indexed read sets for interesting and informative sequences, but recognize that speed and computational resources are an important consideration in the adoption of the method. One obvious, but transient, constraint on the utility of FlexTyper is the ability to generate the FM-index of a read data set. Our implementation utilizes the SDSL library, chosen for its stability, however as the FM-index is critical to many aspects of genome scale analyses there have been strong efforts to develop more efficient indexing algorithms. Recent methods have shown both dramatic increases in construction speed either through induced suffix sorting (Kärkkäinen, Kempa et al. 2017) or GPU-based construction algorithms (Chacón, Marco-Sola et al. 2015), and decreases in memory requirements (Labeit, Shun et al. 2017, Chen, Li et al. 2018). Within our current framework, to try and mitigate some of these issues, we built in methods to split the read set into smaller chunks, each of which is indexed in serial. Although not currently implemented, it is clear that this could be executed in parallel, if there is sufficient RAM available, as each index is generated independently of other chunks. Furthermore, the nature of the reverse mapping approach holds promise with massive parallelization approaches, including those involving GPU acceleration (Hung, Hsu et al. 2018). Moving forward, accelerations to the FM-index generation and reverse mapping approach will result in faster genomic analysis pipelines than is currently possible with alignment-based methods.

Looking to the future, we see the k-mer-searching approach of FlexTyper as having great utility when used in conjunction with emergent pan-genome and graph representations of the reference genome (Kehr, Trappe et al. 2014, Kaye 2016, Paten, Novak et al. 2017). Whether users seek to select a population specific reference graph as the basis for read mapping, or to introduce Bayesian priors (edge weighting) within a pan-population reference graph, knowledge of population markers spanning chromosomes will be required to inform the processes. Furthermore, it is our expectation that pan-genome mapping methods will ultimately use full text read-based indexes, to allow for data compression without loss of information or functionality, while avoiding the plethora of issues facing approaches that index a pan-genomic representation (Paten, Novak et al. 2014, Ghaffaari and Marschall 2019). As the reference structure is enriched and algorithms for use with pan-genome graphs mature, approaches such as FlexTyper, which enable the reverse mapping of informative sequences against a set of indexed reads, will be instrumental in the initial steps of genome analysis pipelines.

