## Supplemental Information for "Demonstrating the utility of flexible sequence queries against indexed short reads with FlexTyper"

For the manuscript:

Richmond & Kaye et al.

**Outline:**

- Supplemental Figures
- Supplemental Tables
- Supplemental Methods
- Supplemental References

### **Supplemental Figures**

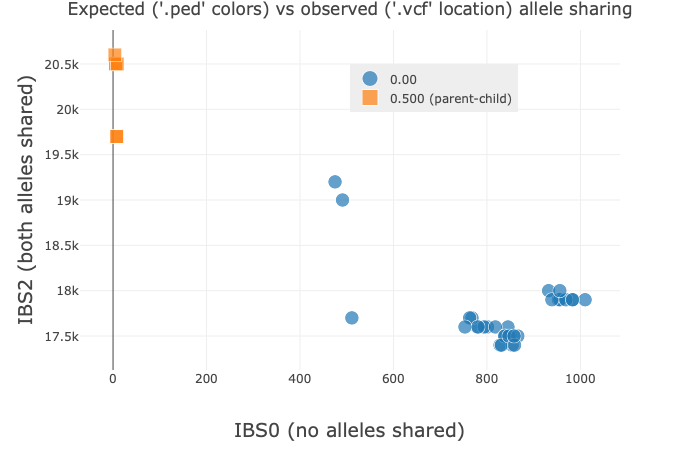

**Figure S1**

For the three trios (nine individuals), the output from Peddy is plotted to inform relatedness. Relatedness comparison for the three trios correctly identifies the six relationships of parent-offspring (orange squares) and correctly identifies a lack of relatedness for the other comparisons.

### **Supplemental Tables**

All Tables are stored in the attached Excel Spreadsheet. Table legends are listed below:

**Table S1**

Indexing times for WGS and RNA seq samples using FlexTyper. Columns include sample id, sample type (WGS or RNAseq) with read pair info (e.g. 2x150), read count, number of sub-indexes, maximum RAM used, wall time, time in seconds. For the WGS samples, we list the time in seconds for the analysis with other tools including BWA mem, Samtools, and DeepVariant.

**Table S2**

An exploration of parameter settings for the recovery of pathogen reads in pure simulated viral samples. Indexed samples include EBV, HIV-1, FR751039, U21941, and ERR2322364 (human RNA-seq sample). The viral samples are simulated to a depth of 100x genomic coverage. Each subsection of the table is the analysis for an individual sample, which has been searched with specified parameters. An example parameter string: k100s1m2000u_UniqRead, can be interpreted as k-mer length 100, step size 1, maximum occurrence 2000, uniqueKmer parameter turned on, and unique read flag turned on. For each of the queries (EBV, HIV-1, FR751039, U21941), the FlexTyper count in Single-end mode is listed. Last, the search time for retrieving k-mers from the index is listed in seconds.

**Table S3**

Extended data for Figure 3, showing the results of the mixed viral analysis with altered parameters for k, s, and unique reads. The sample (Patient 1-5), query strain (EBV, HIV-1, U21941, FR751039, and TOTAL HPV as sum of U21941 and FR751039), expected single end (SE) number of reads, expected paired end (PE) number of read pairs, k-mer length, step size, boolean value for unique read parameter (UniqRead), FlexTyper Single-End count, FlexTyper Paired-End count, Centrifuge paired-end count, and Kraken2 Paired-end count.

**Table S4**

FlexTyper search times for WGS samples. The operation (either Cytoscan search or Ancestry search), number of queries, sample id, max RAM used for search, wall time, and time in seconds. Parameter settings for search found in main text.

### **Supplemental Methods**

##### Data sources for query generation

CytoScanHD SNP probe set was created using the file CytoScanHD_Array.na33.annot.csv acquired from the Affymetrix site (<https://www.thermofisher.com/ca/en/home/life-science/microarray-analysis.html>). This file was processed using CytoscanSProbe2Query.py (<https://github.com/wassermanlab/OpenFlexTyper>).

GRCh37 ancestry sites were acquired from <https://github.com/brentp/peddy/blob/master/peddy/GRCH37.sites>. These sites were then converted into a query file using Sites2Query.py.

Pathogen fasta sequences were acquired from NCBI for EBV (gi|82503188|ref|NC_007605.1| Human gammaherpesvirus 4), HIV-1 (gi|4558520|gb|AF033819.3| HIV-1), U21941.1 (U21941.1 Human papillomavirus type 70), and FR751039.1 (FR751039.1 Human papillomavirus type 68b). Each of these fasta files was then converted into a query file using the fasta_2_query.py script (<https://github.com/wassermanlab/OpenFlexTyper/tree/master/fmformatter>).

African contigs were pulled from NCBI (<https://www.ncbi.nlm.nih.gov/nuccore/PDBU01000000>), and converted into query files using fasta_2_query.py. HLA sequences were downloaded from the IPD-IMGT database (ftp://[ftp.ebi.ac.uk/pub/databases/ipd/imgt/hla/](http://ftp.ebi.ac.uk/pub/databases/ipd/imgt/hla/)), and were parsed to only including the major alleles, based on the allele representation of: GeneName*##:##:##:## where the first ## represents the major allele group  (<http://hla.alleles.org/nomenclature/naming.html>).

##### Samples for genotype analysis testing

To demonstrate the genotyping capacity of FlexTyper at known SNP sites, we used WGS datasets from the Polaris project Diversity Cohort and Kids Cohort (<https://github.com/Illumina/Polaris>). These WGS data sets were downloaded as raw fastqs, mapped against GRCh37 genome using BWA mem (v0.7.5), and then converted to BAM format using Samtools (v1.9). Variants were then called using DeepVariant (v0.10.0) (Li and Durbin 2009, Li, Handsaker et al. 2009). All WGS processing for fastq → vcf  was done on an HPC cluster scheduler using a maximum of 150GB of RAM with 32 CPUs. We additionally tested the indexing capacity using sample HG002, downloaded from <https://ftp-trace.ncbi.nlm.nih.gov/ReferenceSamples/giab/data/AshkenazimTrio/HG002_NA24385_son/NIST_Illumina_2x250bps/reads/>.

##### Simulation of pathogen-containing samples

To simulate pathogen samples in RNA-seq data, we first acquired fasta sequences for four different viruses (described above) and simulated reads at various depths of coverage for paired-end 150bp reads using the read simulator ART (Huang, Li et al. 2012). We tested the recovery of these reads on the simulated files, and additionally spiked them into RNAseq samples. The human RNA-seq samples come from the Genome England project (<https://www.ebi.ac.uk/arrayexpress/experiments/E-MTAB-6523/samples/>). We chose five different human blood RNA-seq datasets to mix with differing read counts of each virus, as defined in Table 1. The patients use blood derived from sample accessions as follows: Patient_1-ERR2322363, Patient_2:ERR2322364, Patient_3:ERR2322365, Patient_4:ERR2322366, Patient_5:ERR2322363.

##### BAM coverage calculation for comparison

Bam coverage is calculated from an input BAM file and a FlexTyper query file using the script QueryFromBam.py (<https://github.com/wassermanlab/OpenFlexTyper/tree/master/extras/bamquery>). This script utilizes the Pysam package to extract the reference and alternate alleles present at a given locus as defined from an input FlexTyper query file.

##### Using Peddy for ancestry, sex, and relatedness typing

After running FlexTyper on the ancestry + chrom X sites, we converted the output format to VCF for input into Peddy. The resulting VCF files for all nine individuals were then compressed with bgzip and indexed with tabix (v1.9), and then merged with bcftools merge (v1.10.1). Next, a PED file was created detailing the relationships and sex of each individual based on information from the Polaris data repository (<https://github.com/Illumina/Polaris>). We used the merged VCF and the PED file as input to Peddy (v0.4.3) and ran with default settings to generate ancestry, relatedness, and sex-typing figures (Pedersen and Quinlan 2017).

##### Running Centrifuge and Kraken2 on simulated patient data

Centrifuge (v1.0.4) was run with default parameters on the simulated “patient” fastqs, where each patient had mixed viral and human blood RNA-seq reads. The output report files from centrifuge were parsed for the viral names using grep: Human immunodeficiency virus 1 (HIV-1), Human herpesvirus 4 type 2 (EBV), and “apilloma” to get all strains of Human (and other) Papilloma viruses (HPV). The numReads (number of reads) were used as a fair comparator to FlexTyper which also was using non-unique read counting for viral detection.

Kraken2 (2.0.9) was run with default parameters on the standard parameters and outputting the mpa-style format, on the default database for human + viral + bacterial genomes (built on June 30th 2020). For collating counts from the mpa-style output report file, with EBV and HIV1 we select the value at the species level as these were accurately assigned. For collating counts from U21941 and FR751039, we extracted counts at the family (Papillomaviridae) level. For subsequent analysis of pure viral reads, we ran Kraken2 independently on 5200 simulated reads from each of the two genomes and collated the output in the same fine.

##### Using FlexTyper: Parameter explanation

**Sliding Search Parameters**

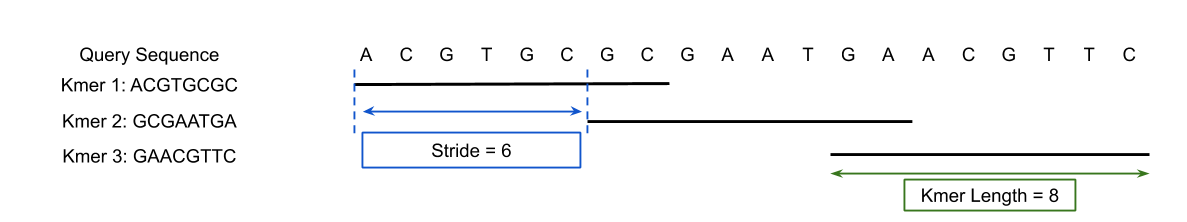

**Centered Search Parameters**

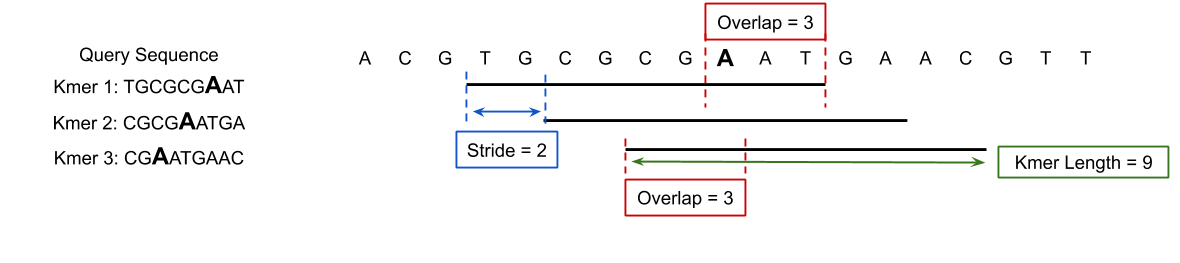

**Non Unique k-mers**

If two query sequences contain an identical k-mer, then that k-mer is flagged as a ‘non unique k-mer’.

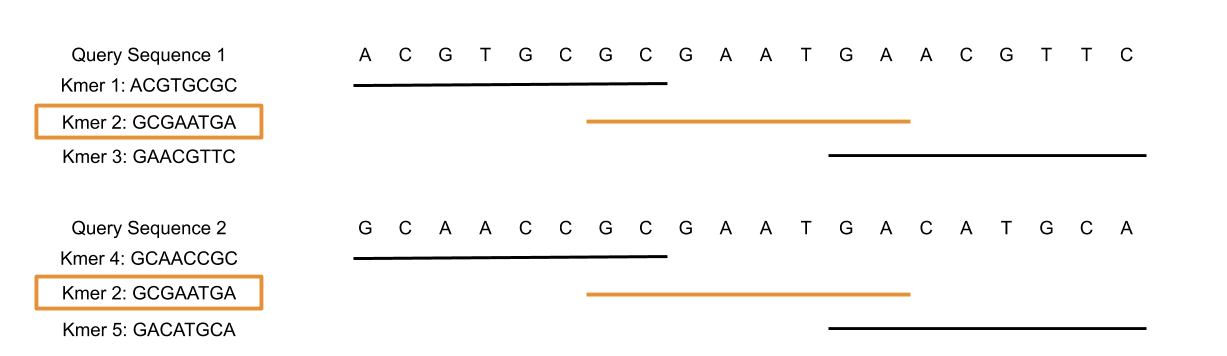

**Over Counted k-mers**

If a k-mer search returns too many positions, then that k-mer is disregarded as uninformative and flagged as an “over counted k-mer”.

**Count As Pairs**

For paired reads [R1 = <1,1>, R2 = <1,2>]

If countAsPairs = true, then if a query matches R1 and R2, then the query count is only one.

**Unique Reads**

For two queries Q1 and Q2, uniqueReads = true, then if a read, R, matches Q1 and Q2, then R will be removed from the set of matching read hits for both Q1 and Q2.

**Combining countAsPairs and uniqueReads**

Suppose you have 3 sets of paired reads:
Pair 1: [R1 = <1,1>, R2 = <1,2>]

Pair 2: [R3 = <2,1>, R4 = <2,2>]

Pair 3: [R5 = <3,1>, R6 = <3,2>]

And two query sequences with matching read hits:
Query 1 = [R1, R3, R5, R6]

Query 2 = [R1, R4]

|  | countAsPairs = True | | | countAsPairs = False | | |
| --- | --- | --- | --- | --- | --- | --- |
|  | Query | Read Hits | Count | Query | Read Hits | Count |
| uniqueReads = True | 1 | [R5, R6] | 1 | 1 | [R3, R5, R6] | 3 |
|  | 2 | [ ] | 0 | 2 | [R4] | 1 |
| uniqueReads = False | 1 | [R1, R3, R5, R6] | 3 | 1 | [R1, R3, R5, R6] | 4 |
|  | 2 | [R1, R4] | 2 | 2 | [R1, R4] | 2 |
